# Resource ratio fluctuations drive the evolution of microbial metabolic strategies

**DOI:** 10.64898/2025.12.18.695187

**Authors:** Zihan Wang, Jacopo Grilli, Akshit Goyal, Sergei Maslov

**Affiliations:** Department of Physics, University of Illinois Urbana-Champaign, Urbana, Illinois, USA; Carl R. Woese Institute for Genomic Biology, University of Illinois Urbana-Champaign, Urbana, Illinois, USA; Quantitative Life Sciences, The Abdus Salam International Center for Theoretical Physics, Trieste, Italy; International Centre for Theoretical Sciences, Tata Institute of Fundamental Research, Bengaluru 560089, India; Department of Bioengineering, University of Illinois Urbana-Champaign, Urbana, Illinois, USA

## Abstract

Microorganisms employ diverse metabolic strategies to cope with environmental variability, yet the evolutionary conditions that favor specialists, hierarchical utilizers, or co-utilizers remain poorly understood. Here, we investigate how fluctuations in resource supply ratios drive the evolution of metabolic strategies. By simulating microbial community assembly in environments with variable resource ratios, we show that diverse metabolic strategies emerge as evolutionary optima in different fluctuation regimes. We find that environments with balanced resource supply select for specialists, whereas imbalanced environments select for generalists. As the magnitude of fluctuations in resource supply ratios progressively increases, generalist strategies smoothly shift from those of hierarchical utilizers to co-utilizers. We identify the duration of temporal niches as the mechanistic driver that selects for reduced lag times over maximal growth rates. These findings demonstrate how the temporal structure of resource availability can spontaneously generate the three distinct metabolic strategies observed in nature.

## Introduction

In natural environments, microorganisms use a variety of metabolic strategies to cope with changes in resource availability. Microbial species are typically able to grow on multiple substrates but not on others [1, 2, 3, 4, 5], displaying a wide range of specialization. Moreover, when multiple resources are provided, they do not always consume them at the same time. In particular, when several resources are present microbes display two broad metabolic strategies: depleting resources one after another (hierarchical utilization) [6, 7, 8] or depleting many available resources simultaneously (co-utilization) [9, 10]. While there is a wide variability across microbial species in this wide spectrum of strategies, we do not understand the conditions in which they can be stably maintained over evolutionary time. If one strategy is always better than the others, we would expect all microbes to adopt it. Hence the diversity of available strategies suggests that different ones might be better at dealing with different environmental conditions.

Very little is known about the evolutionary advantages of different microbial metabolic strategies. Instead, it is better understood how microbes implement these strategies physiologically [11, 12, 13]. Proteome allocation models provide a successful framework to understand possibilities and constraints in expressing metabolic enzymes for utilizing different resources in the environment [14]. In particular, quantitative proteomics studies have revealed that bacteria employ simple strategies for allocating their limited cellular resources across different metabolic functions [15, 16], with partitioning of the proteome fundamentally constraining growth strategies [17].

In the context of growth in the presence of multiple resources, proteome allocation controls both the growth rate of a microbe on currently consumed resources, as well as the lag time when shifting to other resources. Importantly, proteome partitioning creates a natural trade-off between current growth rate and future lag times [18, 19]. For instance, in presence of two sequentially consumed resources, as the fraction of enzymes allocated towards secondary resources decreases, the growth rate on the current primary resource increases, but at the cost of an increased lag time when the resource runs out [19, 20, 21]. The evolutionary plasticity of this allocation should be key to determining which metabolic strategy is optimal in which environment [22, 23], but remains poorly understood, except in the case of a single limiting resource [24, 25, 26]. Further, if resource environments fluctuate, the competition between lag times and fluctuation times may become an important factor driving the evolution of strategies.

Here, using simulations and theory, we demonstrate a simple scenario where all three metabolic strategies — specialists, hierarchical and co-utilizers — can evolve naturally and be maintained stably. We find that allowing the allocation fractions of species to gradually evolve through mutations results in the emergence and stable maintenance of different strategies in different types of fluctuating environments. There are two ways in which resource environment can fluctuate [27, 28]: (1) where overall resource concentrations change over time, e.g., when resources are supplied in discrete chunks. These fluctuations are typically called boom-and-bust environments such as those realized in serial dilution experiments [29, 30]; (2) where there is a imbalance in supplied resource ratios from one boom phase to another [31, 32, 33, 34]. While much of past work has focused on the former kind of fluctations, our work highlights the latter. We suggest that the magnitude of fluctuations in resource supply ratios is the key axis along which different metabolic strategies stratify. We find that specialists are optimal when resource ratios are roughly balanced; hierarchical utilizers when resource ratios have an intermediate imbalance; and co-utilizers when supplied resources ratios are extremely imbalanced. These results show that environmental fluctuations in resource supply ratios may explain how diverse metabolic strategies can stably coexist over long evolutionary time.

## Results

### Modeling evolution of metabolic strategies in fluctuating environments

Our goal is to construct a simple model of microbial community evolution that demonstrates how distinct metabolic strategies can evolve and be maintained in fluctuating environments. To do so, we will make several simplifying assumptions. We assume that evolutionary changes (e.g., through mutations) resulting in changes to proteome allocation are much faster than the ones in metabolic enzymes [35, 36, 37, 38]. Under this assumption, only proteome allocation changes over evolution, while all other parameters (e.g., maximal potential growth rates or lag timescales) stay fixed. We model mutations as continuous changes to proteome allocation, which allows species to transition smoothly from one metabolic strategy to another.

We assume that each microbial species possesses an internal, fixed preference order for the *n*_*R*_ available resources (Methods). The preference order is initialized such that the resource yielding the fastest potential growth rate is assigned rank 1 (“most preferred resource”); the ranking among all other resources is set randomly, reflecting our assumption that there is low evolutionary pressure on the growth rates associated with these lower-ranked choices [39]. Among all *n*_present_ currently available resources, the resource with the highest rank in this order is designated as the primary resource, while the remaining *n*_present_ −1 resources are collectively treated as secondary resources. We parametrize the strategies in the following way: each resource is allocated a fraction *ϕ/n*_present_ of the metabolic proteome; the primary resource is allocated an extra fraction (1 − *ϕ*) to its metabolic proteome.

Thus, the secondary allocation fraction *ϕ* is the key quantity in our model which determines a species’ metabolic strategy (see Methods). Specialists dedicate their entire metabolic portion of the proteome to growth on their sole resource and have no secondary resource; hence for specialists, *ϕ* = 0. Hierarchical utilizers dedicate a dominant fraction of their metabolic proteome to their primary resource and only a small fraction to secondary resources; thus for hierarchical utilizers *ϕ* is nonzero but small. Finally, when the secondary allocation becomes a sizable part of the metabolic proteome, i.e., when *ϕ* becomes comparable to 1, a species’ strategy resembles that of co-utilizers (Fig. 1). Thus, by observing how the secondary allocation fraction *ϕ* evolves in different contexts, we can explore the continuous evolution of metabolic strategies. Importantly, *ϕ* allows to continuously interpolate between these otherwise distinct metabolic strategies.

**Figure 1.**
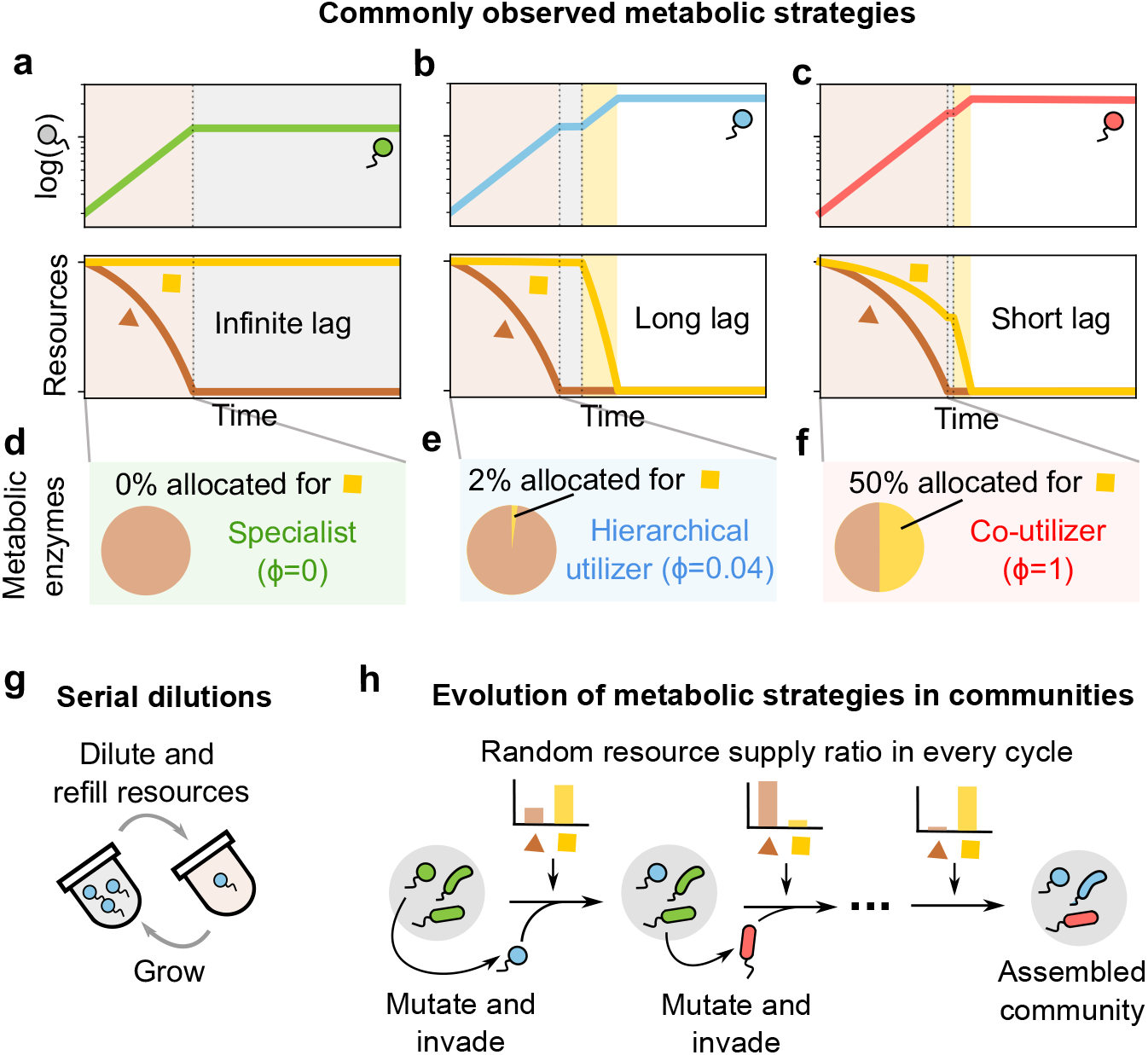
Evolution of metabolic strategies under resource fluctuations. **(a)–(f):** Schematic of three metabolic strategies defined by their secondary allocation *ϕ*. Proteome allocation creates a trade-off: higher *ϕ* shortens the metabolic lag but reduces the initial growth rate. **(a), (d)** A specialist (*ϕ* = 0, green) consumes only the primary resource (brown) and has an infinite switching lag (gray). **(b), (e)** A “hierarchical” utilizer (*ϕ* = 0.05, blue) consumes both resources at the beginning but strongly prioritizes the primary resource. After its depletion, the microbe takes a long lag switching to fully using the secondary resource (yellow). **(c), (f)** A co-utilizer (*ϕ* ≈ 1, red) consumes both resources simultaneously with a minimal switching lag. **(g)–(h):** Simulation of community assembly under fluctuating resources. Microbial communities undergo growth and dilution cycles. In each cycle, a random resource supply ratio is drawn from a Dirichlet distribution (Fig. 2). Starting with a specialist population, new strategies evolve through mutation of *ϕ* and compete, leading to a diverse assembled community.

To model strongly fluctuating environments, we change the resource supply ratios from cycle to cycle. We parameterize these fluctuating resource ratios using a Dirichlet distribution with concentration parameter *α* (Fig. 2a). In each growth cycle, the fraction of total resources supplied as a given resource is drawn from Dir(*α*). As *α* decreases, the distribution of resource ratios shifts from balanced (all resources supplied in roughly equal amounts) to progressively more imbalanced (one or few resources dominate the supply). Thus *α* quantifies the magnitude of resource fluctuations across cycles, with large *α* corresponding to small fluctuations and small *α* to large fluctuations.

**Figure 2.**
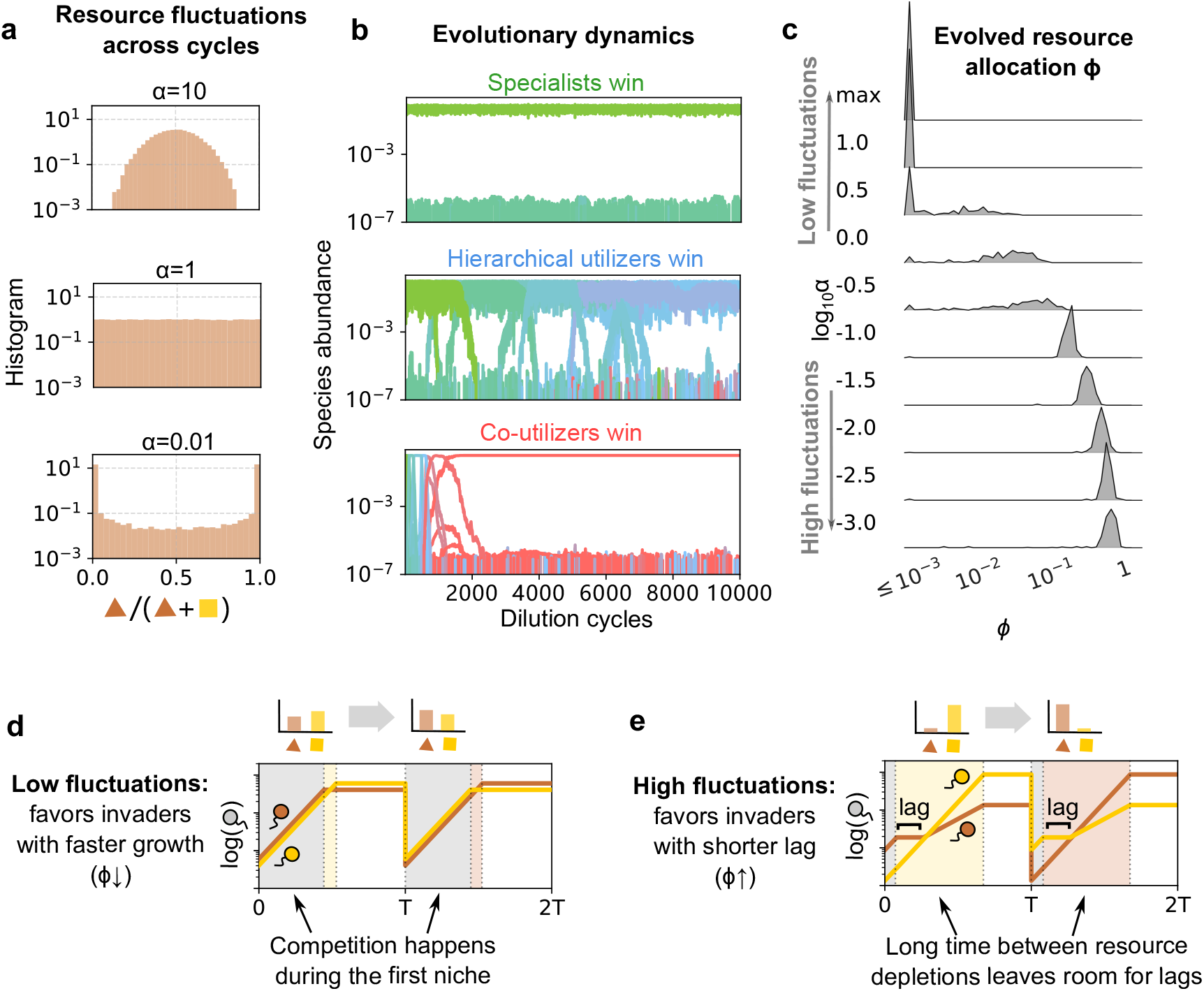
Different metabolic strategies evolve under increasing fluctuations. **(a)** Distribution of resource supply ratios at different strengths of fluctuation. *α* is the concentration parameter from Dirichlet distribution, and is taken at the same value for all resources. The resource supply ratio is balanced (close to 1:1) for large *α* (top), uniformly distributed for intermediate *α* (middle), and fluctuates strongly for small *α* (bottom). **(b)** Example run of community evolutionary dynamics. Colors represent different strategies (green: specialized, blue: hierarchical, red: co-utilizing). At low fluctuations (top), the specialist population is uninvadable. For intermediate fluctuations (middle), final communities consist of hierarchical utilizers, and is uninvadable by co-utilizers. For high fluctuations (bottom), co-utilizers win quickly. **(c)** Ridge plot of secondary allocation *ϕ* in evolved communities. Each row of this plot represents a different fluctuation strength, with the height in each plot representing the frequency of strains with a given *ϕ*—weighted by their abundance—across all evolved communities. log_10_ *α* = max represents equal supply of both resources every cycle (no fluctuations). The lower bound of *ϕ* is clipped to 10^−3^. **(d)–(e)** Different fluctuation strengths select distinct metabolic strategies by changing temporal niche durations. To illustrate this, we consider the coexistence of 2 hierarchical utilizers (*ϕ* = 0.05). Solid lines show strain abundance dynamics across 2 cycles, where colors represent the primary resource of each strain. In each cycle, the first temporal niche (gray) is where both resources are available, and the second temporal niche (light yellow or light brown) is where only 1 resource (corresponding to color) remains. With low fluctuations (d), the secondary niche is much shorter than the first, favoring strains with higher growth rates, and thus lower *ϕ*. With high fluctuations (e), the secondary niche becomes much longer, favoring strains with shorter lags and thus, higher *ϕ*.

### Different metabolic strategies evolve under increasing fluctuations

We simulate community evolution by introducing mutants at a constant rate *µ* = 0.05 per dilution cycle (Methods). Each mutant is generated by selecting a parent individual at random from the community and modifying its secondary allocation fraction by a small amount: log *ϕ*_mutant_ = log *ϕ*_parent_ + Δ log *ϕ*, where the mutation effect size Δ log *ϕ* is drawn from a uniform distribution between −2 and 2. The mutant attempts to invade the resident community. If it can successfully grow and establish itself, it is added to the community; otherwise, it goes extinct. We repeat this process for 10,000 dilution cycles to allow the community to evolve sufficiently.

The results of the evolutionary dynamics reveal a striking pattern across three distinct resource fluctuation regimes (Fig. 2b). At large *α* (balanced resource ratios), specialists with *ϕ* ≈ 0 dominate the evolved communities. At intermediate *α*, we observe the emergence of hierarchical strategies with intermediate secondary allocation fractions of *ϕ* ~ 0.01 −0.1. At low *α* (highly imbalanced resource ratios), co-utilizer strategies with large *ϕ* ~ 0.1 −1 become favored. By collecting statistics from many simulations with randomly chosen parameters, we mapped how the evolved secondary allocation fraction *ϕ* systematically shifts as the magnitude of resource fluctuations increases (Fig. 2c).

### Duration of temporal niches governs evolutionary transitions

Having established that different metabolic strategies emerge in different fluctuation regimes, we now seek to understand the underlying mechanism driving these evolutionary transitions. We first gain intuition by examining a simplified system (Fig. 2d, e) in which a community of 2 resident hierarchical utilizers (*ϕ* = 0.05) with opposite resource preferences has reached a steady state in a periodic environment, where the resource supply ratio alternates between 2 prescribed ratios in successive cycles, rather than fluctuating randomly. The within-cycle dynamics are characterized by distinct temporal niches: the primary niche (Fig. 2d, e, gray), the initial time window where both resources are present, and the secondary niche (Fig. 2d, e, light yellow/brown), the subsequent time window after one resource is depleted, where only the remaining resource is present.

In the low-fluctuation periodic environment (Fig 2d), the system alternates between two ratios that are both close to 1:1. This results in a secondary niche that is too short for either resident species to reallocate their proteome. Competition is thus determined primarily by the growth rate during the long primary niche. A mutant with a lower secondary allocation *ϕ* will be favored and will successfully invade. This is because reducing *ϕ* increases the metabolic proteome dedicated to the primary resource, yielding a faster growth rate during the primary niche, while incurring no penalty for its longer lag time in the secondary niche. Conversely, in the high-fluctuation periodic environment (Fig. 2e), the system alternates between two highly imbalanced ratios. This creates a much longer secondary niche duration. Here, the ability to quickly switch and utilize the remaining resource becomes the critical determinant of fitness. Although increasing *ϕ* reduces the primary growth rate (a disadvantage in the short primary niche), the benefit of a shorter lag time allows the mutant to rapidly exploit the extensive secondary niche. A mutant with a higher *ϕ* (approaching a co-utilizer) will therefore be favored and successfully invade.

In essence, when the primary temporal niche is sufficiently long, competition occurs mainly on the primary resource, favoring rapid growth and small *ϕ* (Fig. 2d), corresponding to specialization. As the primary niche gets shorter compared to the secondary temporal niche, competition instead shifts to secondary resources, favoring short lags for hierarchical utilizers and hence larger *ϕ* (Fig. 2e). The former occurs for low fluctuations (large *α*) while the latter occurs for high fluctuations (small *α*), leading to an increasing optimal *ϕ* with environmental fluctuation strength (Fig. 2c). Although illustrated here in a simplified, periodic steady state, this same fundamental mechanism governs the evolutionary outcomes in the more complex, non-steady state, randomly fluctuating environments investigated in this work.

### Robustness and emergence of three distinct metabolic strategies

While environmental fluctuations do increase the duration of secondary temporal niches, they are not the only way to do so. Indeed, another way is when resources have different intrinsic “qualities”, making some resources systematically better for microbial growth than others [40, 7]. Thus far, we have considered cases when all resources were equivalent and provided similar growth rates. To allow for differences in resource quality, we now introduce a parameter *q. q* = 0 represents no resource quality differences, which was the case for Fig. 2. As *q* increases, so does the systematic difference between growth rates on different resources (Methods).

The effect of resource quality is most clearly illustrated by considering a community of two specialists in a non-fluctuating environment (*α* = max). When resource quality is equal (Fig. 3a), resources are depleted roughly simultaneously, creating a primary temporal niche (gray) and virtually no secondary niche. This state is evolutionarily stable; any mutant with *ϕ >* 0 would be outcompeted by the specialists’ faster growth rate in the primary niche and would gain no advantage, as there is no secondary niche to exploit. However, when one resource has a higher quality (e.g., *q* = 0.3, Fig. 3b), it is always depleted first, if provided in equal supply. This in itself creates a stable secondary temporal niche (light yellow) where only the lower-quality resource (yellow) remains. This newly created niche favors the emergence of hierarchical mutants preferring the higher-quality resource (brown) to invade. By adopting a hierarchical strategy (*ϕ >* 0), such mutants grows slower on the primary resource but crucially, they compensate for this disadvantage by having a short enough lag which allows them to consume the lower-quality resource during the secondary niche before it gets depleted by other species.

**Figure 3.**
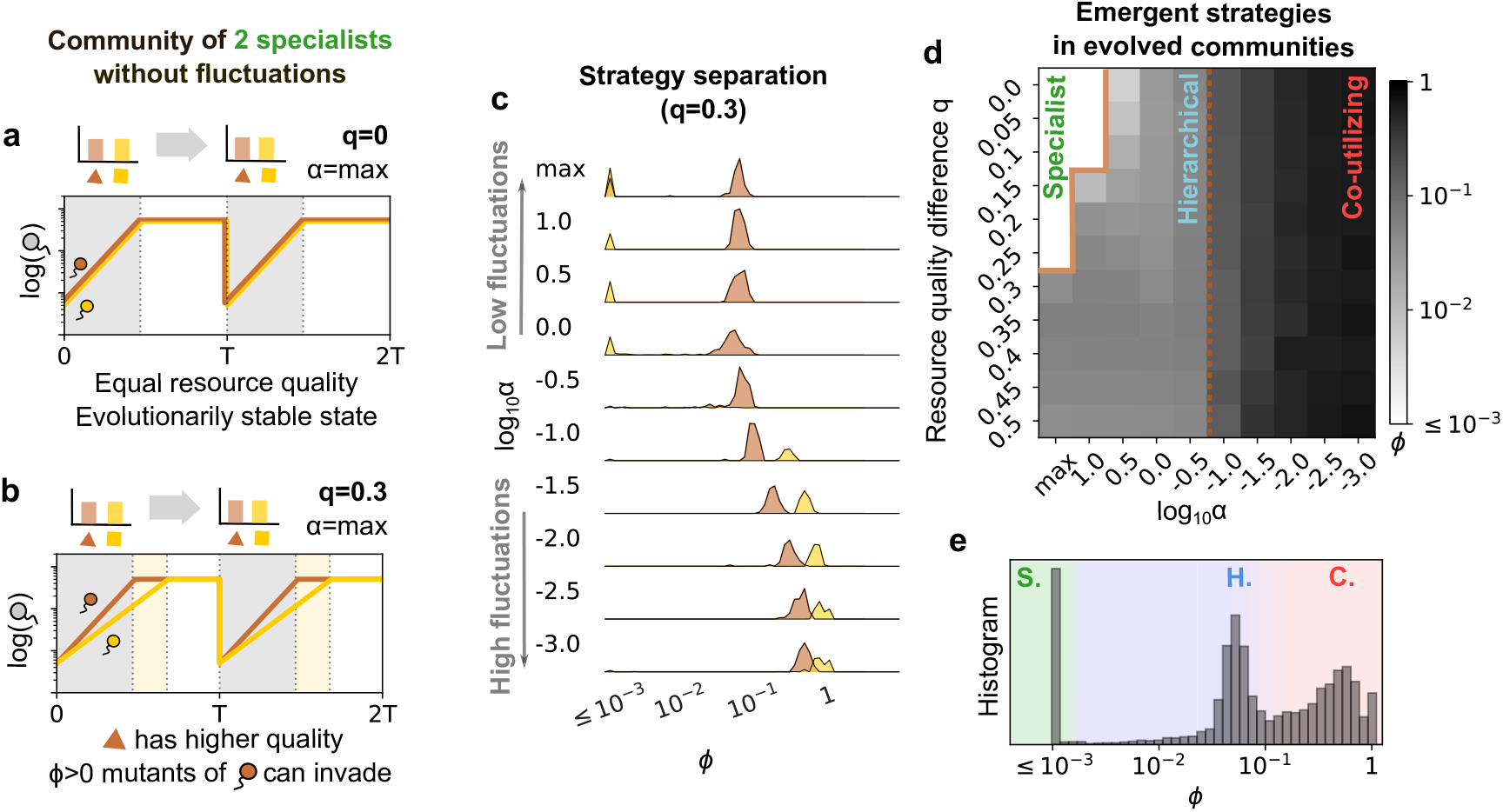
Emergence of three distinct metabolic strategies. **(a)–(b)** Resource quality difference *q* affects temporal niches when there is no fluctuation. We consider the coexistence of 2 specialists (*ϕ* = 0). Solid lines show the abundance change across 2 cycles, where colors represent the primary resource of each species. (a) When both resources have the same quality (*q* = 0), they will be depleted simultaneously (gray), and 2 specialists will reach an uninvadable state to mutations in *ϕ*. (b) when the brown resource has higher quality (*q* = 0.3), the yellow resource will always be depleted later, creating a second temporal niche (light yellow). Mutants of the brown species whose *ϕ >* 0 will therefore be able to invade. **(c)** Ridge plot of secondary allocation *ϕ* in final communities, where colors (brown and yellow) imply the primary resource of the corresponding species. At each fluctuation strength, the y-axis stands for the relative abundance of species of a given *ϕ* across all communities. log_10_ *α* = max represents no fluctuation. The lower bound of *ϕ* is clipped to 10^−3^. **(d)** Emergent strategies from simulated evolutionary dynamics in each environment. x-axis represents the resource supply ratio fluctuation, and y-axis stands for the resource quality difference. Grayscale from white (*ϕ* ≤ 0.001) to black (*ϕ* = 1) stands for averaged *ϕ* across multiple simulation runs. The solid brown line separates specialists and hierarchical utilizers at *ϕ* = 0.001, and the dotted brown line separates hierarchical with co-utilizing species at *ϕ* = 0.1. **(e)** Histogram of *ϕ* for all evolved strains across all environments in (d), weighted by their relative abundances. 3 distinct peaks emerge, corresponding to 3 metabolic strategies: specialist (S), hierarchical (H), and co-utilizing (C).

We therefore expect that evolution favors a specific path for the emergence of hierarchical utilizers. This is precisely what we observe in our simulations. At *q* = 0.3 (Fig. 3c), strains preferring the higher-quality resource (brown) contribute the majority of mutants which become hierarchical utilizers (intermediate *ϕ*). The species preferring the lower-quality resource (yellow) largely remains specialized (*ϕ* ≤ 0.001) until environmental fluctuations become very high.

By repeating our evolutionary simulations, similar to those in Fig. 2, for different values of *q*, we plotted the average value of *ϕ* in evolved communities (Fig. 3d). This plot clearly shows that even when resources have different quality, increasing the degree of environmental fluctuations (decreasing *α*) systematically shifts the evolved strategies from specialists to hierarchical to co-utilizing strategies. Once resources become sufficiently different in their quality (*q* ~ 0.3), specialists are displaced by hierarchical utilizers, which dominate over a wider range of environmental fluctuations *α*. Pooling all evolved communities across the full range of *q* and *α* revealed that the secondary resource allocation *ϕ* followed a strikingly trimodal distribution (Fig. 3e). The three peaks of this distribution allowed us to partition regions of *ϕ* into three discrete metabolic strategies: specialists (S, green) at *ϕ* ≤ 0.001, hierarchical utilizers (H, blue) between 0.001 < *ϕ* < 0.1 and co-utilizers (C, red) between 0.1 < *ϕ* < 1. Thus, across a variety of environments — with different resource qualities and fluctuations — the process of evolution naturally produces three clearly distinguishable metabolic strategies.

In our simulations so far, we have evolved communities supplied with two resources. We find that our central results also hold when communities are evolved in environments with more resources *n*_*R*_ = 3 and *n*_*R*_ = 4 (Fig. 4). Specifically, across several environments with varying resource quality difference *q* and environmental fluctuation strength *α*, we continue to observe a trimodal distribution of evolved secondary allocation *ϕ*, corresponding to specialists (S), hierarchical utilizers (H) and co-utilizers (C) (Fig. 4a,d). Further, with increasing fluctuation imbalance *α*, the evolved communities transition from mostly comprising specialists at large *α*, hierarchical utilizers at intermediate *α* and finally co-utilizers at low *α* (Fig. 4b,e). This demonstrates the robustness of our central result. Finally, while the maximal species diversity equals the number of resources *n*_*R*_ due to competitive exclusion [41, 42], we observe maximally diverse communities for the largest values of *α* (balanced resource supply). These environments typically support communities with *n*_*R*_ specialists, each specializing on a different resource (Fig. 4c,f). As *α* decreases, environmental fluctuations increase, and community diversity drops. At low *α* values, evolved communities are largely dominated by 1 co-utilizer. For intermediate *α* values, communities contain specialists and hierarchical utilizers, which are typically complementary on their most-preferred resource (Supp. Fig. 4) [39].

**Figure 4.**
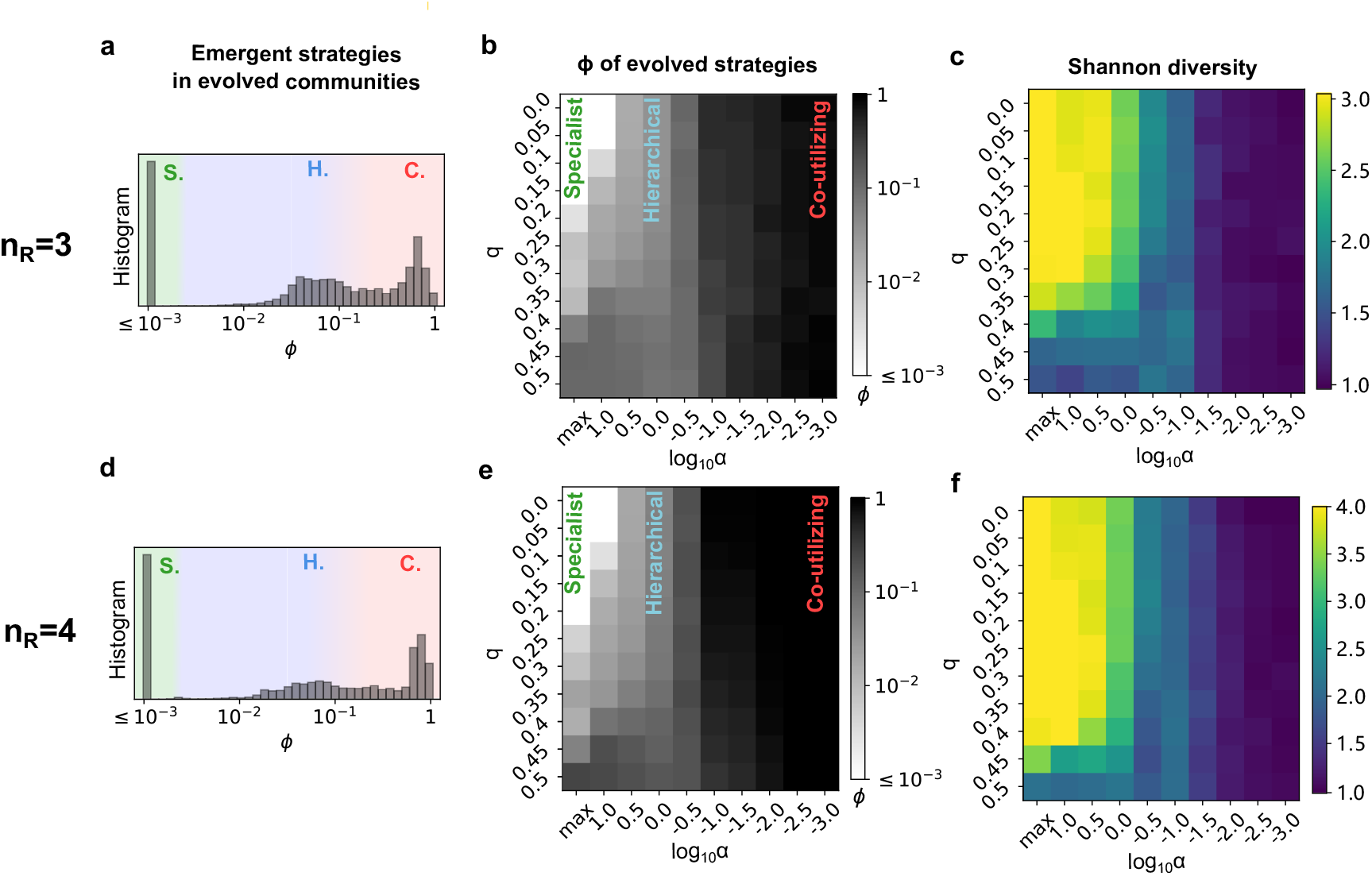
Emergence of three distinct metabolic strategies is robust to increasing number of resources. **(a), (d)** Histograms of *ϕ* for evolved strains show the emergence of specialists (S), hierarchical utilizers (H), and co-utilizers (C), similar to Fig. 3d. **(b), (e)** Heatmaps of average *ϕ* at each environment shows that the evolved metabolic strategies of *n*_*R*_ = 3, 4 share a similar pattern with *n*_*R*_ = 2 as shown in Fig. 3c. **(c), (f)** Heatmap of average effective number of species (*e*^*H*^ where *H* is Shannon diversity) in different environments. x-axis represents the strength of resource supply ratio fluctuations, and y-axis stands for the resource quality difference. Colorbar represents a range from 1 to *n*_*R*_. Specialists and hierarchical utilizers tend to form more diverse communities, whereas co-utilizing communities, evolved under high resource ratio fluctuations, are usually dominated by one strain.

## Discussion

In recent years, there has been a substantial focus on understanding microbial community assembly and dynamics in boom-and-bust environments typified by serial dilution experiments [39, 7, 34, 30, 23]. Virtually all these studies focus on changing parameters such as the dilution factor, representing the severity of the “bust” phase of dynamics, as well as the time between dilutions and the total supplied resource concentration, which represents the duration and strength of the “boom” phase respectively. However, to our knowledge, with the exception of Ref. [7], all these studies maintain a balanced resource ratio which remains constant from one growth cycle to the next. While this makes experiments simple and tractable, it is far from realistic for natural environments, where resources are typically supplied in ratios that fluctuate from cycle to cycle [28, 43]. In this study, we demonstrate a remarkable consequence of these fluctuations in resource ratio imbalance. We find that changing the magnitude of resource supply fluctuations promotes the evolution of distinct metabolic strategies: from specialists dominating at balanced ratios (low fluctuation magnitude) to hierarchical utilizers at intermediate magnitude, to co-utilizers dominating in environments with highly imbalanced resource supply ratios. These results potentially explain the diversity of metabolic strategies observed in natural microbial communities.

The mechanism driving these evolutionary transitions is rooted in the temporal structure of resource depletion within each growth cycle. Balanced resource ratios create only brief secondary temporal niches (after the first resource depletion), while imbalanced ratios generate extend the duration of these secondary niches. Specialists and hierarchical utilizers prioritize rapid growth on their primary resource and consequently have long switching lags—an optimal strategy when secondary niches are too short to exploit. Co-utilizers sacrifice some growth rate on the primary resource to gain the ability to grow during extended secondary niches. Hierarchical utilizers occupy an intermediate position, maintaining sufficient primary growth while developing short enough lag times to capitalize on meaningful secondary niches. This mechanistic understanding explains why increasing environmental fluctuations does not universally favor generalist strategies, but rather selects for the strategy whose proteome allocation best matches the temporal niche structure imposed by that fluctuation regime.

Our findings are robust across resource quality differences and numbers of available resources. Resources with different qualities promote secondary temporal niches even in non-fluctuating environments through systematic differences in their depletion time. Yet the fundamental pattern persists: increasing imbalance in resource supply ratios systematically shifts community composition from specialists to hierarchical utilizers to co-utilizers. Surveying evolved species across environments, we consistently observe a trimodal distribution of secondary allocation fractions corresponding to three distinct metabolic strategies.

Our model is intentionally minimal and makes several simplifying assumptions. First, we assume that evolutionary changes occur primarily through proteome reallocation (mutations adjust allocation fractions) on timescales faster than the evolution of the metabolic machinery itself, so maximal growth rates are held fixed while allocation evolves continuously. In addition, each strain has a fixed resource-preference order that does not evolve over these timescales. These assumptions, while simplifying, are supported by experiments showing that allocation (and hence lag times) can evolve quite readily, even when maximal enzyme-catalytic rates or resource preference hierarchies show little to no change even after thousands of generation of evolution [44, 45]. Future work that allows several of these parameters to co-evolve could further our understanding of how metabolic strategies emerge and stably coexist.

These findings have direct implications for understanding natural microbial diversity. One testable prediction of our work would be that microbial communities isolated from environments with increasingly imbalanced resource supply ratios would have a progressively increasing ratio of co-utilizing to hierarchically utilizing or specialist species. Even though it is a relatively straightforward experimental task to infer the metabolic strategy of an isolated microbial species [6, 7], to our knowledge, it is rather challenging to infer the extent of imbalance of resource ratio fluctuations in natural environments.

## Acknowledgements

We thank G. Chure for valuable discussions. This research was supported in part by the International Centre for Theoretical Sciences (ICTS) during a visit for participating in the program — Winter School on Quantitative Systems Biology 2025 (Code: ICTS/Prog-qsb/2025/1). A.G. acknowledges support from the Ashok and Gita Vaish Junior Researcher Award, the Govt. of India’s Ramanujan Fellowship, as well as the DAE, Govt. of India, under project no. RTI4001. J.G. acknowledges the support of Fondo Italiano per la Scienza - FIS (CUP J53C23002290001). The research of Z. W and S.M. was supported in part by grants from the NSF (DMS-2235451) and Simons Foundation (MPS-NITMB-00005320) to the NSF-Simons National Institute for Theory and Mathematics in Biology (NITMB).

## Data and code availability

There are no data associated with this paper. All code is available as a GitHub repository at the following link: https://github.com/maslov-group/Evolution of metabolic strategies

## Methods

### Tradeoff between growth rate and lag time mediated by secondary allocation *ϕ*

For simplicity in modeling, we assume that the metabolic sector of the proteome allocation is constitutive throughout the growth phase. At any given time point during growth, a species *α* partition its metabolic proteome across multiple resources. This allocation is represented by fractions *ϕ*_*αk*_, which is the fraction of the metabolic proteome allocated to consume resource *k*, subject to the constraint ∑_*k*_ *ϕ*_*αk*_ = 1.

In our model with *n*_*R*_ resources, we assign each species a preference order for all *n*_*R*_ resources. When *n*_present_ resources are present, each species will take the one with the highest rank on its preference order as the primary resource. For each species *α*, the top of its preference order (“most preferred resource”) will be resource *k* that provides the fastest growth rate max_*k*_(*g*_*αk*_); the rest *n*_*R*_ − 1 resources will be ranked randomly [39]. We assume that the proteome fraction allocated to any single secondary resource will not exceed the fraction allocated to the primary resource. Based on this assumption, the allocation to each present secondary resource is defined as:

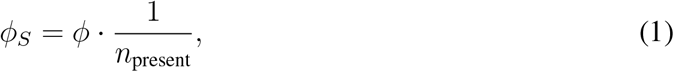

where *ϕ* is the scaled secondary allocation, constrained between [0, 1]. *ϕ* = 0 corresponds to specialists that do not allocate to secondary resources, and *ϕ* = 1 corresponds to the idealized co-utilization strategy where the total metabolic proteome is allocated equally among all *n*_present_ available resources. Consequently, the allocation to the primary resource is:

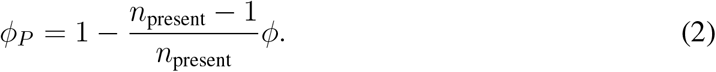

And the allocation to any absent resource is set to zero. As a result, the overall growth rate of species *α* is given by the weighted sum of potential growth rates: *g*_*α*_ = ∑_*k*_ *g*_*αk*_*ϕ*_*αk*_. As the scaling parameter *ϕ* increases, the proteome allocation to the primary resource (*ϕ*_*P*_) decreases, leading to a corresponding reduction in the overall growth rate (*g*_*α*_).

When a resource *k* is depleted, species that were previously consuming it need to reallocate their proteome to focus on the remaining present resources. Such reallocation of proteome, results in a gradual increase of growth rate, effectively corresponding to a lag-time. We model this as a dynamic process where the existing enzymes for all present resources are used to synthesize new metabolic enzymes specifically for the available resources. Denote *ϕ*_present_ = 1 − *ϕ*_depleted_ as the fraction of such enzymes, we then model the rate of change of this fraction as *dϕ*_present_*/dt* = *ϕ*_present_*/τ*_0_, where *τ*_0_ is a timescale. The lag phase duration, *τ*, is defined as the time required for the allocated fraction *ϕ*_present_ to increase from its initial value (at the moment of depletion) up to 1. Integrating the above differential equation from *t* = 0 to *t* = *τ*, we obtain the expression for the lag phase duration:

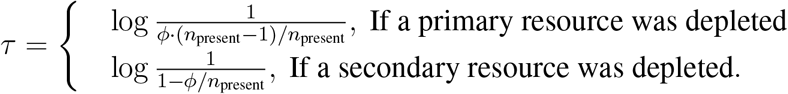

This lag is reduced as *ϕ* increases.

### Model of community dynamics in boom-and-bust environments

We model the dynamics of a microbial community operating in a boom-and-bust environment. The system consists of *n*_*R*_ substitutable resources, with concentrations *R*_*k*_ (*k* = 1, 2, …, *n*_*R*_), and *n*_*S*_ strains, with abundances *N*_*α*_ (*α* = 1, 2, …, *n*_*S*_).

The environment is characterized by a periodic cycle of period *T*. At the beginning of each cycle, resources are supplied in large bulks; at the end of each cycle, all that is left in the system is diluted by a factor of *D*. The cycle period *T* is assumed to be much longer than the inverse of the maximum growth rates (*g*_*α*_*T* ≫ *D*), ensuring that all resources are completely depleted within every cycle.

We assume that the initial resource concentrations are significantly higher than the species’ half-saturation constants. As a result, species grow exponentially at constant rates (*g*_*α*_) during the growth phase. The consumer-resource dynamics are described by the following set of equations:

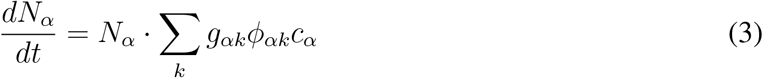

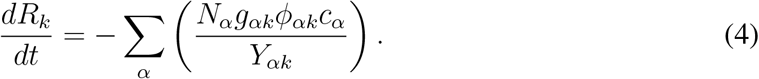

Here, *Y*_*αk*_ is the yield, which we set to 1 for all pairs of species and resources for the sake of simplicity. *c*_*α*_ is a binary switch for species *α*: it is set to 0 if all resources are depleted or *α* is in lag phase, and to 1 if *α* is during active growth.

### Details of numerical simulations

In all simulations, the maximum potential growth rate *g*_*αk*_ of species *α* on resource *k* is calculated in the following way:

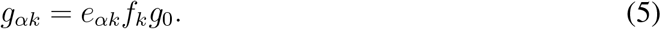

Here, *g*_0_ = 0.8 hr^−1^ represents the upper bound (or maximal potential) growth rate. The term *e*_*αk*_ is a coefficient that quantifies species *α*’s intrinsic ability to utilize resource *k*. The coefficient *f*_*k*_ represents the quality of resource *k*.

To model the difference in resource quality, denoted by the parameter *q*, across *n*_*R*_ resources, we simplify the structure by ranking the resources from 1 (best) to *n*_*R*_ (worst). The resource quality factors *f*_*k*_ are then defined by the relation: 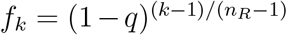. This design ensures that the best resource (*k* = 1) has a quality factor *f*_1_ = 1, and the worst resource (*k* = *n*_*R*_) has 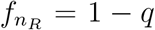. We restrict the quality difference parameter to the range *q* ∈ [0, 0.5] for our simulations.

Drawing inspiration from our previous work in serial dilutions [39], where selective pressure is often strongest on the utilization of the most preferred resource, and little on other resources, we assign a “most preferred resource” to each species *α*. For the species’ most preferred resource, the utilization coefficient is set to *e*_*αk*_ = 1. For all other resources, the coefficient is set to *e*_*αk*_ = *p*. In our simulations, we fix *p* = 0.5. Given that *q* ∈ [0, 0.5], we have 1 − *q* ≥ 0.5 = *p*. This ensures that the resource designated as the “most preferred” for any species is also the one yielding the absolute fastest *g*_*αk*_ for that species.

We set the lag timescale at *τ*_0_ = 0.3 hr. It is important to note that the actual lag times (*τ*) experienced by our hierarchical utilizers are typically significantly longer than the characteristic timescale *τ*_0_. The serial dilution parameters are set as follows: the period of a single cycle is *T* = 24 hours, and the dilution factor is *D* = 100.

In our numerical simulations of randomly fluctuating resource ratios, the resource supply vector at the beginning of each dilution cycle is sampled independently from the Dirichlet distribution Dir(*α*, …, *α*).

Each simulation run is initialized with a community of *n*_*R*_ specialist species (where *ϕ* = 0), with each species specializing in a different resource. In every cycle, there is a probability *µ* = that an existing species mutates. The mutant species acquires a new proteome allocation parameter *ϕ*^*′*^ and invades the community. For the initial specialist species (*ϕ* = 0), the only possible mutation is to *ϕ*^*′*^ = 10^−4^. For species with non-zero *ϕ*, the new parameter is calculated as *ϕ*^*′*^ = *ϕ·*exp(*s*), where the mutation step *s* is drawn from a uniform distribution *s* ~ *U* (−2, 2). The resulting *ϕ*^*′*^ is then clipped to the interval [10^−4^, 1]. We do not consider mutations in preference orders. Every cycle, there is a probability *p*_migrate_ = 0.005 that a random specialist species from the initial community will invade.

All invading species (mutants and migrants) start with a low initial abundance of 10^−8^. Furthermore, any existing species whose abundance falls below 10^−8^ at the beginning of a cycle is considered extinct and is removed from the simulation. For each specific set of environmental conditions, we perform 1000 independent simulation runs, with each run encompassing 10, 000 serial dilution cycles. We take the last 10 cycles of every run and combine them to create the dataset of evolutionary outcomes.

## Notes

https://github.com/maslov-group/Evolution_of_metabolic_strategies

### Competing Interest Statement

The authors have declared no competing interest.

