## Supplementary Information for "Resource ratio fluctuations drive the evolution of microbial metabolic strategies"

This file contains:

Supplementary Text

Supplementary Figures 1-7

### Supplementary Text

#### Simplified periodic resource supply protocol

To analytically investigate the mechanisms observed in the numerical experiments, we replace the random resource supply protocol with a periodic one. This simplification allows for a steady-state analysis of resource competition and generalist invasion. We consider the simplest scenario involving $n_R = 2$ resources ($R_1$ and $R_2$). The resource supply fluctuation is defined over two sequential dilution cycles:

1. Odd cycle (Cycle $I$): Resource supply is $[R_1, R_2] = [R_L, R_H]$.
2. Even cycle (Cycle $II$): Resource supply is $[R_1, R_2] = [R_H, R_L]$.

This preserves the mean resource availability and fluctuation magnitude observed in the original numerical experiments characterized by the Dirichlet concentration parameter $\alpha$. The lower and higher resource supply amounts, $R_L$ and $R_H$, are calculated as the expected minimum and maximum, respectively, from the Beta distribution (Supp. Fig. 1a):

$$R_L(\alpha) = E[\min(R_1, 1 - R_1)] = \frac{2 \cdot B_{0.5}(\alpha + 1, \alpha)}{B(\alpha, \alpha)} \tag{S.1}$$

and $R_H(\alpha) = 1 - R_L(\alpha)$. Here, $B(x, y)$ is the Beta function.

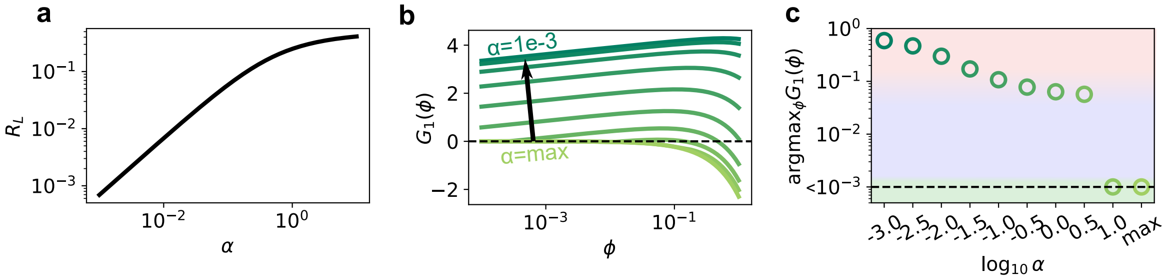


Supplementary Figure 1: **Invasion of generalists to the specialist community is affected by environmental fluctuation strength.** **(a)** Converting random resource supply to periodic. A random environment characterized by $\alpha$ is converted to a periodic environment represented by $[R_L,\ 1 - R_L]$. **(b)** Net growth of a generalist when it invades a community of 2 specialists. The invasion is possible only if $G_1(\phi) > 0$. x-axis marks the $\phi$ of the invader generalist, and y-axis represents its net exponential growth across 2 cycles. Colors represent the fluctuation strengths: from low fluctuation ($\alpha = \max$, lime) to high fluctuation ($\alpha = 10^{-3}$, turquoise), as in (c). **(c)** Optimal strategy and first-order phase transition. Points show the optimal invading strategy ($\phi^* \equiv \mathrm{argmax}_\phi G_1(\phi)$) as a function of the environmental fluctuation $\alpha$. $\phi^*$ is the allocation strategy $\phi$ that yields the maximum net growth for the corresponding colored curve in panel (b). All values lower than $10^{-3}$ are plotted at $10^{-3}$ for visualization. Colors in the background indicate strategies: co-utilizing (red), hierarchical (blue), and specialist (green).

#### First-order phase transition between specialists and generalists

We first analyze the steady-state coexistence of two specialist strains, $N_1$ and $N_2$. Strain $N_1$ only consumes $R_1$, and strain $N_2$ only consumes $R_2$. The resource-dependent growth rates are defined as $g_{\alpha k}$, the growth rate of species $\alpha$ on resource $k$. Following the assumptions of equal resource quality ($q = 0$), we set the maximum growth rates as $g_{11} = g_{22} = g_0$, and growth potentials on non-preferred resources as $g_{12} = g_{21} = pg_0$ (see Methods).

In a stable steady state, both specialists must exhibit a net growth factor of $D^2$ across the two-cycle period (where $D$ is the dilution factor). Let us denote that $N_1$ grows by a factor of $\gamma_1$ during the first cycle and by a factor of $\gamma_2$ during the second cycle. We then have $\gamma_1\gamma_2 = D^2$. By symmetry, $N_2$ grows by $\gamma_2$ during the first cycle, and $\gamma_1$ during the second cycle. Mass conservation then gives the following:

$$N_1(0)(\gamma_1 - 1) = R_L \tag{S.2}$$

$$N_1(0)\gamma_1(\gamma_2 - 1)/D = R_H \tag{S.3}$$

$$N_2(0)(\gamma_2 - 1) = R_H \tag{S.4}$$

$$N_2(0)\gamma_2(\gamma_1 - 1)/D = R_L \tag{S.5}$$

Solving these coupled equations for the growth factors yields:

$$\gamma_1 = \frac{R_H + DR_L}{R_L + DR_H}D, \quad \gamma_2 = \frac{R_L + DR_H}{R_H + DR_L}D \tag{S.6}$$

The resource depletion times for the specialists in each cycle are determined by their respective growth factors and growth rates:

| | **Cycle** $I$ | **Cycle** $II$ |
|---|---|---|
| $R_1$ Depletion Time ($T_1$) | $T_1^{(I)} = \frac{\log\gamma_1}{g_{11}}$ | $T_1^{(II)} = \frac{\log\gamma_2}{g_{11}}$ |
| $R_2$ Depletion Time ($T_2$) | $T_2^{(I)} = \frac{\log\gamma_2}{g_{22}}$ | $T_2^{(II)} = \frac{\log\gamma_1}{g_{22}}$ |

Given our growth rate assumptions ($g_{11} = g_{22} = g_0$), the resource ordering will be $T_1^{(I)} < T_2^{(I)}$ and $T_1^{(II)} > T_2^{(II)}$, meaning $R_1$ is depleted first in Cycle I, and $R_2$ is depleted first in Cycle II.

We then consider the invasion of a generalist mutant $N_1'$, derived from $N_1$. $N_1'$ maintains the same single-resource growth rates but gains an allocation factor $\phi$, which allows it to consume $R_2$. The criterion for successful invasion is that the total net exponential growth $G_1(\phi)$ over the two-cycle period must be positive.

At the time of invasion, the biomass of the generalist is minuscule, so it barely affects the depletion times. The net exponential growth is then calculated by summing the growth during the two resources' co-existence phase and the subsequent single-resource phase in each cycle, including lag time:

$$\begin{aligned} G_1(\phi) = {} & [(1-\phi/2)g_{11} + (\phi/2)g_{12}]\,(T_1^{(I)} + T_2^{(II)}) \\ & + g_{12}(\Delta T_1 - \tau_0\log(2/\phi))\Theta(\Delta T_1 - \tau_0\log(2/\phi)) \\ & + g_{11}(\Delta T_2 + \tau_0\log(1-\phi/2))\Theta(\Delta T_2 + \tau_0\log(1-\phi/2)) \\ & - \log D^2 \end{aligned} \tag{S.7}$$

where $\Delta T_1 = T_2^{(I)} - T_1^{(I)}$, $\Delta T_2 = T_1^{(II)} - T_2^{(II)}$, and $\Theta(x)$ is the Heaviside theta function. The extra growth phase for the generalist mutant, during which only one resource remains, is calculated using the lag time equations from the Methods section, where $\tau_0$ is the characteristic lag time scale.

Using the physiological and serial dilution parameters set from Methods, we can plot the net growth $G_1(\phi)$ across the allocation strategy $\phi$ (Supp. Fig. 1b). We observe that the specialist community is uninvadable in environments with weak fluctuations (large $\alpha$). However, as fluctuation strength increases (smaller $\alpha$), the optimal strategy switches abruptly ($\mathrm{argmax}_\phi G_1(\phi) \sim 0.1$), indicating a first-order phase transition where generalist mutants successfully invade (Supp. Fig. 1c).

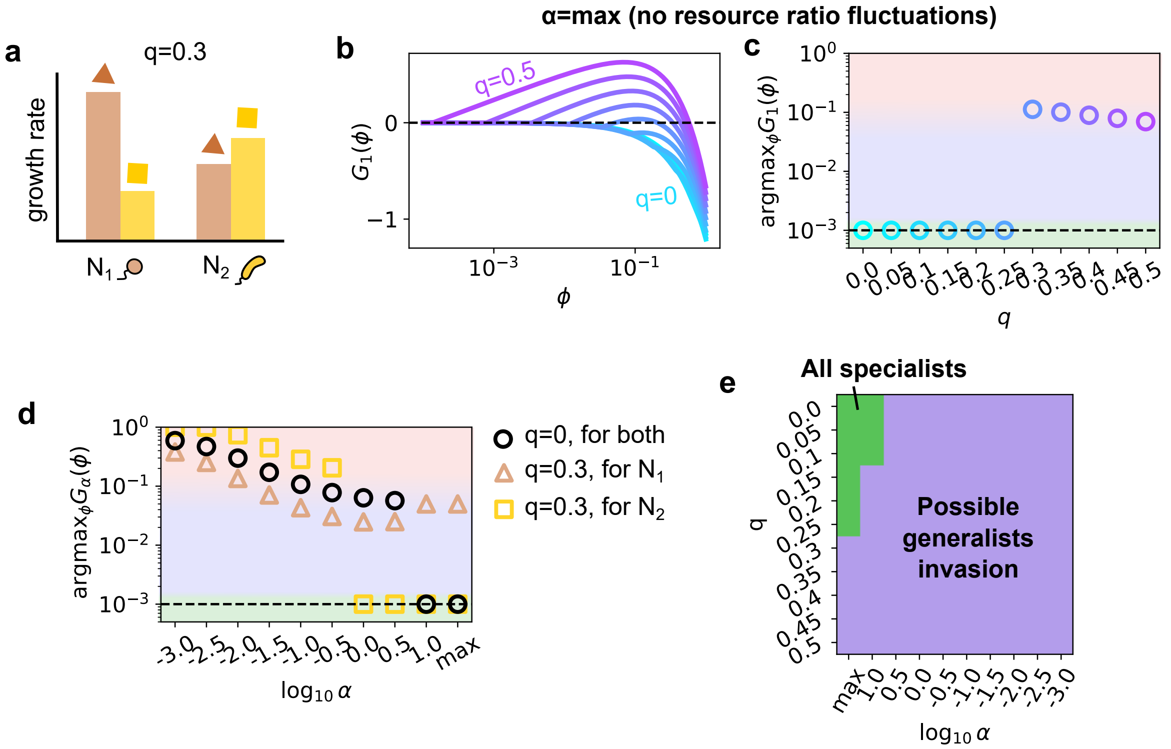


Supplementary Figure 2: **Invasion of generalists to the specialist community is affected by resource quality difference.** **(a)** Example of $q = 0.3$. Growth rates on resource 2 (yellow square) are reduced compared to resource 1 (brown triangle). **(b)** Net growth of a generalist mutant of $N_1$ when it invades a community of 2 specialists. The invasion is possible only if $G_1(\phi) > 0$. x-axis marks the $\phi$ of the invader generalist, and y-axis represents its net exponential growth across 2 cycles. With the resource supply ratio being fixed, colors represent the resource quality difference: from $q = 0$ (cyan) to $q = 0.5$ (purple), as in panel (c). **(c)** Optimal strategy and first-order phase transition. Points show the optimal invading strategy ($\phi^* \equiv \text{argmax}_\phi G_1(\phi)$) as a function of the resource quality difference $q$. $\phi^*$ is the allocation strategy $\phi$ that yields the maximum net growth rate for the corresponding colored curve in panel (b). All values lower than $10^{-3}$ are plotted at $10^{-3}$ for visualization purpose. Colors in the background indicate strategies: co-utilizing (red), hierarchical (blue), and specialist (green). **(d)** Comparing the $\alpha - \phi^*$ relationship in different $q$. Black circles: same as in Supp. Fig. 1c, where $q = 0$. Brown triangles: optimal strategies for $N_1$'s mutants when $q = 0.3$. Note that generalists can always invade regardless of the environmental fluctuations. Yellow squares: optimal strategies for $N_2$'s mutants when $q = 0.3$. The fluctuation region for generalists to invade shrinks compared to $q = 0$. **(e)** Specialist community's invadability across different environments.

We now break the resource symmetry by introducing the resource quality difference $q$. We assume $R_2$ is the resource of lower quality, such that the growth rate on $R_2$ is reduced for both species (Supp. Fig. 2a):

$$g_{22} = (1-q)g_{11} = (1-q)g_0$$
$$g_{12} = (1-q)g_{21} = (1-q)pg_0$$

The growth rates on $R_1$ remain $g_{11} = g_0$ and $g_{21} = pg_0$. Note that the depletion times $T_2^{(I)}$ and $T_2^{(II)}$ are now inversely dependent on $(1-q)$.

This discrepancy in maximum growth rates can break the resource depletion order constraint $T_1^{(II)} > T_2^{(II)}$ established in the symmetric case. For example, in the environment with no supply ratio fluctuation ($\alpha \to \infty$, where $R_H = R_L = 1/2$ and $\gamma_1 = \gamma_2 = D$), if $q > 0$, $R_1$ will always be depleted earlier than $R_2$. The specialist $N_1$, consuming $R_1$, thus gains a strong fitness motive to acquire the generalist trait $\phi$ to consume $R_2$ during the post-depletion phase of $R_1$. In general, a nonzero $q$ encourages $N_1$ to mutate towards generalists but discourages $N_2$ from doing so.

The net exponential growth of mutants, $G_1(\phi)$ and $G_2(\phi)$, must now account for all possible scenarios of resource depletion ordering within Cycle II, as the sign of $T_1^{(II)} - T_2^{(II)}$ is now dependent on $q$. They can be written down as follows:

$$\begin{aligned} G_1(\phi) = {} & [(1-\phi/2)g_{11} + (\phi/2)g_{12}]\,[T_1^{(I)} + \min(T_1^{(II)}, T_2^{(II)})] \\ & + g_{12}(\Delta T_1 - \tau_0 \log(2/\phi))\Theta(\Delta T_1 - \tau_0 \log(2/\phi)) \\ & + g_{11}(\Delta T_2 + \tau_0 \log(1-\phi/2))\Theta(\Delta T_2 + \tau_0 \log(1-\phi/2))\Theta(\Delta T_2) \\ & + g_{12}(-\Delta T_2 - \tau_0 \log(2/\phi))\Theta(-\Delta T_2 - \tau_0 \log(2/\phi))\Theta(-\Delta T_2) \\ & - \log D^2, \end{aligned} \tag{S.8}$$

and

$$\begin{aligned} G_2(\phi) = {} & [(1-\phi/2)g_{22} + (\phi/2)g_{21}]\,[T_1^{(I)} + \min(T_1^{(II)}, T_2^{(II)})] \\ & + g_{22}(\Delta T_1 + \tau_0 \log(1-\phi/2))\Theta(\Delta T_1 + \tau_0 \log(1-\phi/2)) \\ & + g_{21}(\Delta T_2 - \tau_0 \log(2/\phi))\Theta(\Delta T_2 - \tau_0 \log(2/\phi))\Theta(\Delta T_2) \\ & + g_{22}(-\Delta T_2 + \tau_0 \log(1-\phi/2))\Theta(-\Delta T_2 + \tau_0 \log(1-\phi/2))\Theta(-\Delta T_2) \\ & - \log D^2 \end{aligned} \tag{S.9}$$

If we take the environment with no supply ratio fluctuation ($\alpha \to \infty$) and plot how $G_1(\phi)$ changes with $\phi$, we see that for small $q$ values the specialists are uninvadable. However, as $q$ increases, a first-order phase transition happens and some hierarchical utilizers will invade (Supp. Fig. 2b-c).

Furthermore, if we fix $q = 0.3$ and compare the mutants of $N_1$ and $N_2$, we find that generalist mutants of $N_1$ are able to invade specialists ($G_1(\phi) > 0$) for any $\alpha$. Conversely, $N_2$'s mutants tend to stay specialists more often than in the $q = 0$ case (Supp. Fig. 2d), which is consistent with observations in random environments (Fig. 3c). This analytical approach allows us to test all environments, reproducing the specialist uninvadability region found in the main text (Supp. Fig. 2e, Fig. 3d).

#### Supplementary Figures

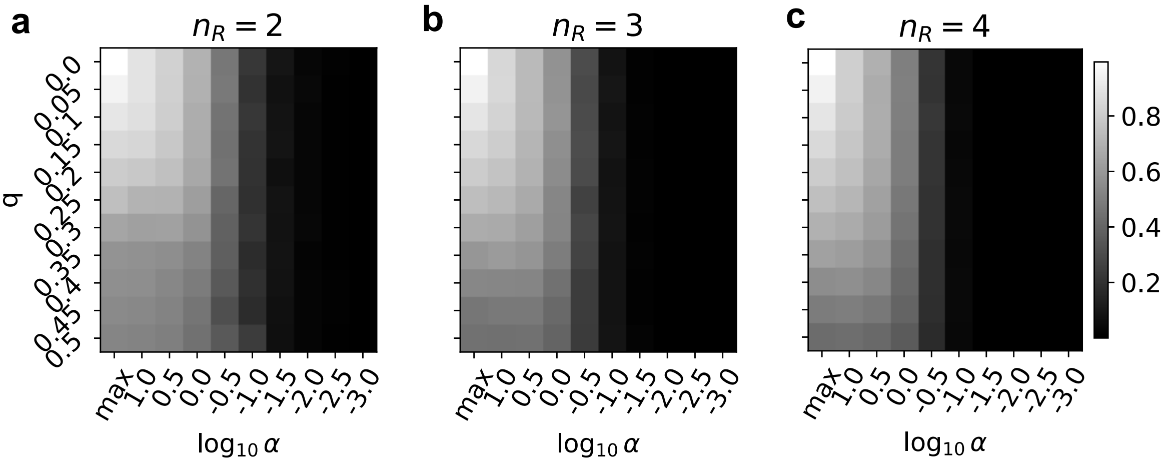


Supplementary Figure 3: **Fraction of growth time during the first temporal niche.** Heatmaps shows the average fraction of total growth time spent when all resources are present in different environmental conditions. Panels (a)-(c) respectively show results for $n_R$=2, 3, 4 resources.

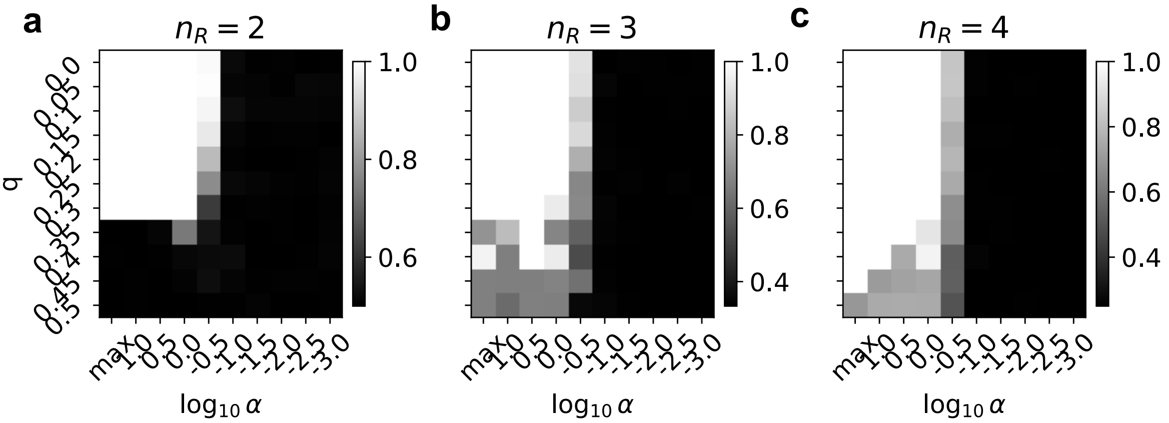


Supplementary Figure 4: **Complementarity of the most preferred resource.** Heatmaps shows the average top choice complementarity, which is calculated as a number of unique resources among the most preferred choices of all species in the community, divided by the number of unique resources $n_R$, ranging between $[1/n_R, 1]$. Panels (a)-(c) respectively show results for $n_R$=2, 3, 4 resources.

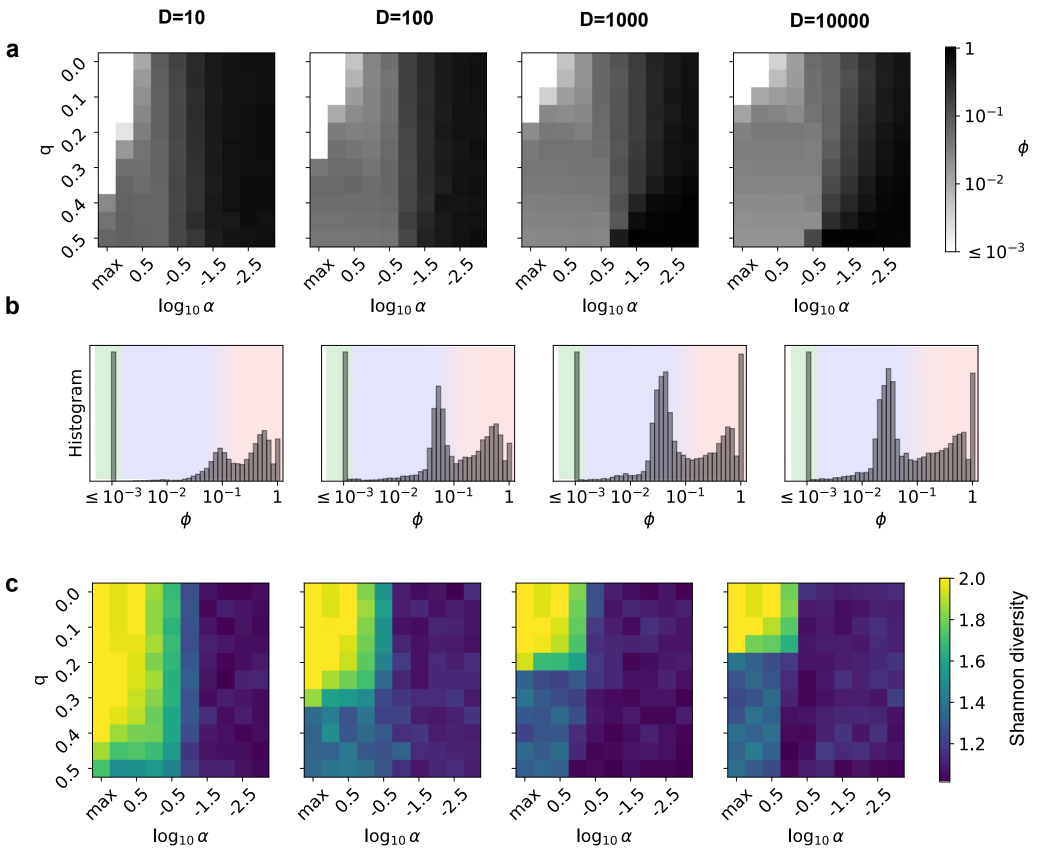


Supplementary Figure 5: **Effects of the dilution factor** $D$**.** All parameters except for $D$ are taken at the same values as in Fig. 3. **(a)** Heatmaps of averaged $\phi$ (represented by grayscale) at each environment across fluctuations (x-axis) and resource quality difference (y-axis). **(b)** Histograms of final evolved strategies ($\phi$) show the emergence of specialists (green), hierarchical utilizers (blue), and co-utilizers (red). **(c)** Heatmap of averaged Shannon diversity at different environments.

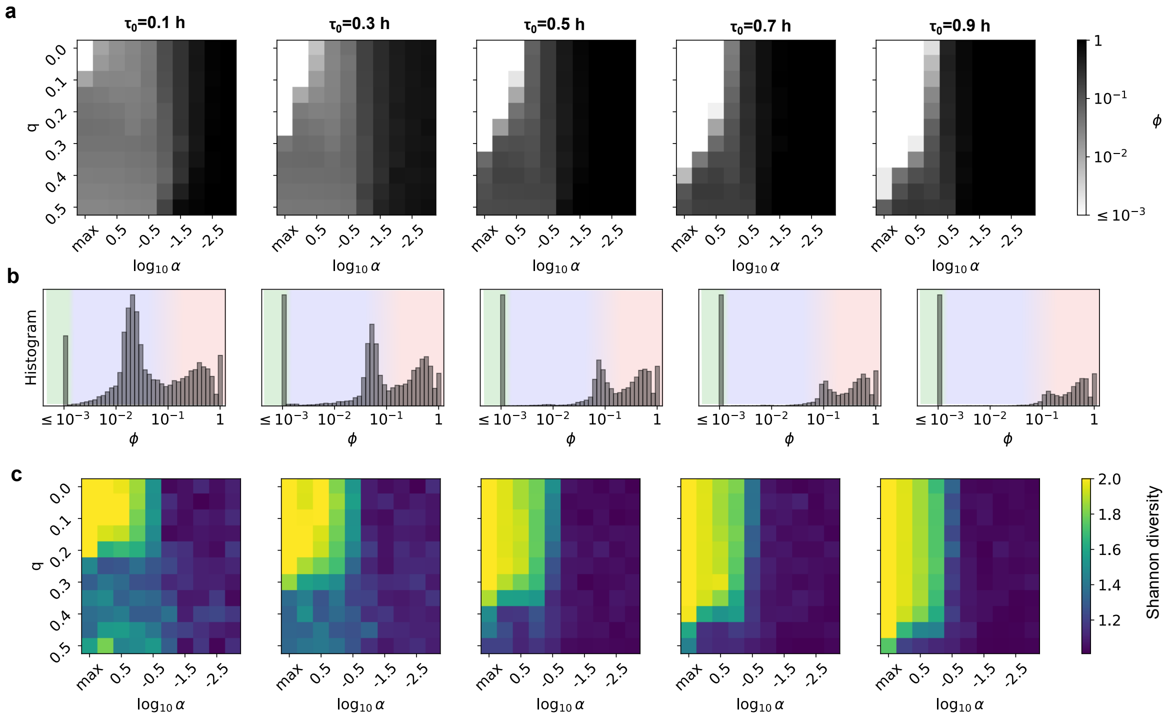


Supplementary Figure 6: **Effects of the lag time scale $\tau_0$.** All parameters except for $\tau_0$ are taken at the same values as in Fig. 3. **(a)** Heatmaps of averaged $\phi$ (represented by grayscale) at each environment across fluctuations (x-axis) and resource quality difference (y-axis). **(b)** Histograms of final evolved strategies ($\phi$) show the emergence of specialists (green), hierarchical utilizers (blue), and co-utilizers (red). **(c)** Heatmap of averaged Shannon diversity at different environments.

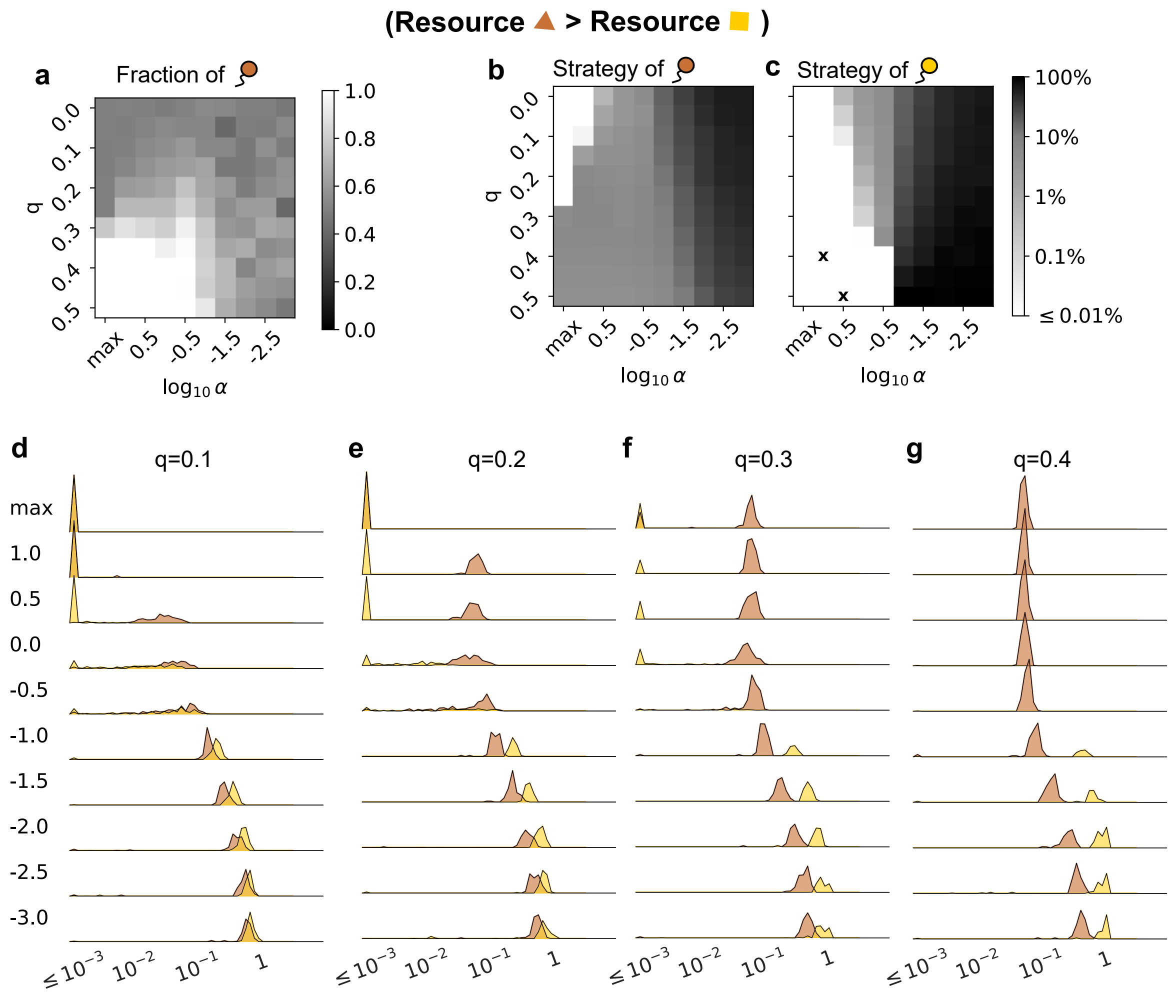


Supplementary Figure 7: **Impact of resource quality difference on evolutionary outcomes.** All parameters are taken at the same values as in Fig. 3. When $q > 0$, the brown resource always has a higher quality than the yellow resource. **(a)** Abundance heatmap of species primarily using the brown resource. Grayscale shows the fraction of such species in total biomass. At $q = 0$ the fraction would be symmetric ($\approx 0.5$). The white region in the lower-left corner shows that strains of the brown species dominate the community when resource fluctuation is low and the quality difference is high. **(b)-(c)** Evolved metabolic strategies. The grayscale represents averaged final $\phi$ value for the species in the final community whose primary resource is brown (panel b) or yellow (panel c) respectively. As $q$ increases, strains of the brown species tend to evolve towards being hierarchical utilizers (gray), and strains of the yellow species often evolve towards specialists (white) or co-utilizers (black) depending on the environmental fluctuation. The "x" marks in panel (c) denote conditions where the strains of yellow species were absent in all simulated outcomes. **(d)-(g)** Ridge plot of secondary allocation $\phi$ in final communities at different $q$ values, where colors (brown and yellow) imply the primary resource of the corresponding species.
